# Major host transitions are modulated through transcriptome-wide reprograming events in *Schistocephalus solidus*, a threespine stickleback parasite

**DOI:** 10.1101/072769

**Authors:** François Olivier Hébert, Stephan Grambauer, Iain Barber, Christian R Landry, Nadia Aubin-Horth

## Abstract

Parasites with complex life cycles have developed numerous phenotypic strategies, closely associated with developmental events, to enable the exploitation of different ecological niches and facilitate transmission between hosts. How these environmental shifts are regulated from a metabolic and physiological standpoint, however, still remain to be fully elucidated. We examined the transcriptomic response of *Schistocephalus solidus*, a trophically-transmitted parasite with a complex life cycle, over the course of its development in an intermediate host, the threespine stickleback, and the final avian host. Results from our differential gene expression analysis show major reprogramming events among developmental stages. The final host stage is characterized by a strong activation of reproductive pathways and redox homeostasis. The attainment of infectivity in the fish intermediate host – which precedes sexual maturation in the final host and is associated with host behaviour changes – is marked by transcription of genes involved in neural pathways and sensory perception. Our results suggest that un-annotated and *S. solidus*-specific genes could play a determinant role in host-parasite molecular interactions required to complete the parasite’s life cycle. Our results permit future comparative analyses to help disentangle species-specific patterns of infection from conserved mechanisms, ultimately leading to a better understanding of the molecular control and evolution of complex life cycles.

## INTRODUCTION

Parasites with multiple hosts commonly undergo dramatic phenotypic transformations and endure major environmental shifts over the course of their life cycle (Wilbur 1980; Poulin 2011), yet very little is known about how these are orchestrated at the molecular and physiological levels, or how conserved they are across species (Auld & Tinsley 2014). Among the key insights yet to be gained is a detailed understanding of the metabolic and developmental regulation of parasites associated with infection, survival and development in each host. Characterising patterns of gene expression can inform the study of how physiological functions are modulated, a task otherwise difficult to achieve for organisms such as parasites that need to be cultured and studied inside other animals. Gathering this information for multiple host-parasite systems will allow general comparisons to be drawn between species. These comparisons will ultimately help disentangle species-specific patterns from common mechanisms that have promoted the evolution of complex life cycles.

We dissected the genome-wide transcriptional activity of the cestode *Schistocephalus solidus*, a model parasite with a complex life cycle (Barber 2013), to uncover how biological functions are regulated in different developmental stages, and how they relate to the completion of the parasite’s life cycle (Figure 1a). *S. solidus* successively parasitizes a cyclopoid copepod, a fish – the threespine stickleback, *Gasterosteus aculeatus –* and a piscivorous endotherm, typically a bird (Clarke 1954). We aimed to determine the functional changes happening when infecting the final host – where reproduction occurs – and identify differences in gene expression between pre-infective and infective forms of the plerocercoid stage within the second intermediate host. The first developmental stages – free swimming coracidia – occur in freshwater and hatch from eggs released with the faeces of the avian host. Coracidium ingested by cyclopoid copepods may then develop further into the procercoid stage. When a threespine stickleback feeds on infected copepods, the parasite is released from the copepod and penetrates the intestinal mucosal wall of the fish after 14-24 hours, before developing into the plerocercoid stage (Hammerschmidt & Kurtz 2007). However, the newly developed plerocercoid is not initially infective to the final bird host. The status of infectivity is defined as the development stage at which the parasite can successfully mature and reproduce (Tierney & Crompton 1992). During the early plerocercoid phase, the host immune system is not activated by the presence of the worm inside its body cavity (Scharsack *et al*. 2007). *S. solidus* spends the next 50-60 days in an exponential growth phase, gaining up to 10 000 times its initial mass (Scharsack et al. 2007). When the plerocercoid eventually reaches infectivity, a phase that could be determined by sufficient glycogen reserves (Hopkins 1950), drastic phenotypic changes occur in the fish host (Barber & Scharsack 2010). These changes include an activation of the immune system (Scharsack *et al*. 2007) and a loss of anti-predator response (Barber *et al*. 2004). Following the ingestion of infected fish by an avian predator, the parasite experiences a temperature of 40°C in the bird’s digestive tract – compared to a maximum of 15-18°C in the ectothermic intermediate hosts – as well as chemical attack by digestive enzymes. These conditions trigger the parasite’s development to the sexually mature adult in ca. 36 hours (Smyth 1950). The adult parasite reproduces during the next 3-4 days with eggs being released into the water with the avian host’s faeces (Hopkins & Smyth 1951). Adjusting to these host switches and life history transitions requires many physiological changes that are expected to recruit the activity of different genes at each phase.

**Figure 1.**
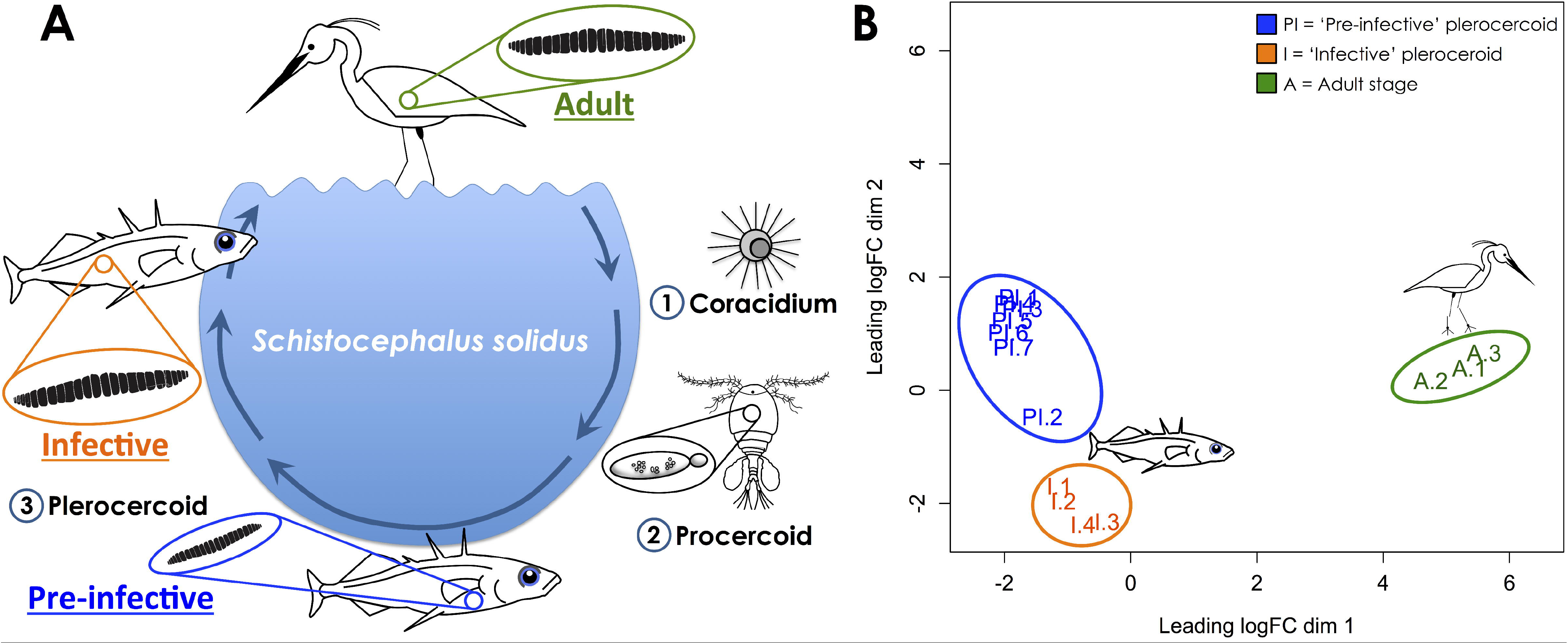
Developmental stages of *S. solidus* are characterised by different genome-wide expression profiles. A) Life cycle of *Schistocephalus solidus.* B) Multidimensional scaling analysis (MDS) confirming the presence of three distinct phenotypes among samples (n=17). The distance between two given points on the graph corresponds to the typical log2-fold-change between the two samples for the top 1000 genes with the largest Euclidian distance.

One of the major transitions expected to affect worm physiology is the transition from somatic growth to reproduction. Histological and physiological studies suggest that gametogenesis only occurs when the parasite reaches the final (bird) host (Hopkins 1950; Schjørring 2003). Despite the advanced (‘progenetic’; Smyth & MacManus 2007) development of reproductive organs in infective plerocercoids, only an elevated temperature of 40°C in semi-anaerobic conditions can trigger meiosis and reproductive behaviours (Smyth 1952; Schjørring 2003). Previous work on the anaerobic activity of key enzymes involved in the catabolism of carbohydrates in *S. solidus* also suggests that while carbohydrate breakdown is very slow in pre-infective and infective plerocercoids, this rate increases several-fold upon maturation (Körting & Barrett 1977; Beis & Barrett 1979). Energetic resources used during the adult stage mainly come from glycogen reserves accumulated during growth of the pre-infective plerocercoid (Hopkins 1952; Körting & Barrett 1977). One thus expects that plerocercoids cultured at 40°C will show an up-regulation of glycogen-related pathways.

The adult stage interacts with its environment to time these developmental steps. The anatomical structure more likely to achieve this task is the tegument, a very active and complex tissue that behaves like a true epidermis (Lee 1967). The adult stage of *S. solidus* exhibits numerous vacuolate vesicles packed with electron-dense or electron-lucent content. These small structures are evenly distributed in *S. solidus* syncytial tegument (Charles & Orr 1968). Their role could be related to both nutrition and defence, as they allow rapid internalization of environmental nutrients and antigen-antibody complexes (Hopkins *et al*. 1978; Threadgold & Hopkins 1981). However, the uptake of macromolecules by adult cestodes remains an open question (Conradt & Peters 1989). If, however, *S. solidus* performs pinocytosis or endocytosis at any developmental stage, specific transcripts involved in this biological activity are expected to be up-regulated at these stages. The existence of membrane-bound vesicles also suggests potential secretory/excretory functions that would allow the parasite to release various types of molecules in its host.

Pre-infective and infective plerocercoids are discrete developmental stages distinguished by their divergent growth and effects on the host immune system. Parasites grow rapidly in the first weeks of the stickleback host infection but growth rates tend to slow down as the parasite becomes infective to the final host (Barber & Svensson 2003). Concurrently, empirical evidence shows that secretory/excretory products from pre-infective versus infective plerocercoids have different modulatory effects on the immune system of the fish host (Scharsack *et al*. 2013). Small, pre-infective plerocercoids down-regulate the proliferation of host monocytes, but as soon as they attain infectivity they activate a strong respiratory burst activity (Scharsack *et al*. 2007). From a transcriptional perspective, different functional programs between pre-infective and infective stages that reflect these divergent activities should be detectable. Distinct and specific gene expression profiles should characterize each developmental stage according to the biological activities that they need to perform to ultimately maximize the parasite’s success in each host.

## MATERIAL AND METHODS

### Sampling

Worm specimens spanning three development states were extracted from laboratory-raised and experimentally infected threespine sticklebacks (detailed protocol in Hébert et al. 2016a). To infect these fish, we obtained parasite eggs through *in vitro* culture of mature plerocercoids extracted from wild-caught threespine sticklebacks (Clatworthy Reservoir, England, UK). After a three-week incubation period in tap water, eggs hatched in response to daylight exposure, and emergent coracidia were used to infect copepods. Laboratory cultured copepods (*Cyclops strenuus abyssorum*) were individually exposed to one coracidium larva. Exposed copepods were screened after three to four weeks for the presence of a single parasite inside its body cavity under a binocular microscope. Exposed copepods harboring infective procercoids were fed to healthy lab-bred threespine sticklebacks (Hébert *et al*. 2016b). One hundred fish had food withheld for 48h, in order to increase the likelihood of fish feeding on infected copepods, and were then individually isolated in 1L plastic tanks. Each fish was fed a single infected copepod and left in its individual tank for one week, with water change every 48h. Exposed fish were subsequently divided into five groups of 20 individuals. Each group was placed in a 50L temperature-controlled aquarium and fed frozen chironomid larvae *ad libitum* for 17 weeks. Exposed fish were randomly selected and euthanized in a benzocaine solution (15 mM) between 10 and 17 weeks post-exposure. The timing of the sampling was determined using known growth curves for the parasite at this water temperature. Our infection protocol resulted in single infections only (one worm / fish body cavity). Pre-infective plerocercoids (i.e. <50 mg) were collected between 10 and 13 weeks post-infection (n = 7) and infective plerocercoids (i.e. >50mg) were collected between 16 and 17 weeks post-infection (n = 4). We obtained three additional adult specimens of *S. solidus* through *in vitro* culture of infective plerocercoids extracted from wild-caught sticklebacks of the same population as the experimental infections (Hébert *et al*. 2016a). These wild fish had spent ten months in the laboratory prior to worm extraction. Therefore, we expected that the parasites extracted from these fish would be comparable with individuals of earlier stages that resulted from experimental infections. We used this approach since we obtained only a 15% infection rate with our experimental infections and needed more *Schistocephalus* individuals than we could collect. Obtaining adult worms through *in vitro* culture in a bird-gut model represents a standard method used for more than 50 years in experimental parasitology applied to helminths. It offers quick and replicable sampling, as compared to alternative methods such as *in vivo* infections of ducklings. Briefly, adult worms were collected after five days of *in vitro* culture in a bird-gut model at 40°C. To do so, infective plerocercoids were placed individually into a dialysis membrane suspended in a medium composed of 50:50 RPMLhorse serum, at a temperature, pH and oxygen tension mimicking the conditions experienced in the bird digestive track (Smyth 1950) – for detailed protocol see (Hébert *et al*. 2016a). Adult worms had a body mass of 321-356 mg. Worms were washed with ultra-pure RNase-free water, diced into small pieces of ~5 mm x 5 mm, placed into RNALater (Ambion Inc., Austin, TX, USA) and kept at -80°C.

### RNA sequencing

We used RNA samples from fourteen different worms to produce individual TruSeq Illumina sequencing libraries (San Diego, CA, USA) according to the manufacturer’s protocol. We produced libraries for seven pre-infective (<50mg) plerocercoids, four infective plerocercoids (>50mg) and three adult worms (Hébert *et al*. 2016). cDNA libraries were sequenced on a Illumina HiSeq 2000 system (Centre de Recherche du CHU de Québec, Québec, QC, Canada) with the paired-end technology (2X100 bp). In total, 75.8 Gb of raw data was generated, which represents 375 million 2 x 100 bp paired-end sequences distributed across the fourteen samples – deposited into the NCBI Sequence Read Archive (accession number SAMN04296611, BioProject PRJNA304161, see Hébert *et al*. 2016c).

### Short-read alignment on reference transcriptome

Raw sequencing reads were cleaned, trimmed and aligned on the reference transcriptome, allowing the estimation of transcript-specific expression levels for each individual worm (Hébert *et al*. 2016b; Hébert 2016b). In summary, we aligned short reads from the 14 individual HiSeq libraries on the reference with Bowtie 2 v.2.1.0 (Langmead & Salzberg 2012), allowing multimapping of each read. Transcripts showing similar sequence, length and expression levels were then regrouped into clusters of unigenes by using Corset v1.00 (Davidson & Oshlack 2014). We obtained read counts for each unigene in each individual worm using the mapping information contained in the SAM files (Hébert 2016b). We adjusted the algorithm parameters so that read counts with sequences of varying lengths (isoforms, pseudogenes, alternative transcripts, paralogs) would not be merged (contig ratio test parameter switched on). The reference transcriptome was annotated with functional information for each gene and published as an open access resource with downloadable reference files (Hebert et al. 2016b). We used this functional annotation for downstream analyses presented in this study.

### Differential expression analysis

We conducted downstream analyses using the R packages ‘limma-voom’ (Law *et al*. 2014) and edgeR (Robinson *et al*. 2010). We imported the read count matrix into R v.3.3.2 (R Development Core Team, 2008) for initial filtering of lowly expressed transcripts. We kept sequences with more than 15 Counts Per Million (CPM) in at least three samples in any of the life stage. The filtering threshold values were chosen based on a comparative analysis of multiple datasets produced with different combinations of thresholds (Figure S1). This specific threshold allowed filtering low-coverage transcripts and potentially several false positives, without losing too much information on differentially expressed transcripts. Normalization of read counts was performed using the method of Trimmed Mean of M-values (TMM), using edgeR default parameters, followed by the voom transformation. Read counts were converted into CPM value and log2-transformed. Next, each transcript was fitted to an independent linear model using the log2(CPM) values as the response variable and the group – pre-infective plerocercoid, infective plerocercoid and adult – as the explanatory variable. No intercept was used and all possible comparisons between the three developmental stages were defined as our desired contrasts. Each linear model was analyzed through limma’s Bayes pipeline. This last step allowed the discovery of differentially expressed transcripts based on a False Discovery Rate (FDR) < 0.001 (Hébert 2016a; Law *et al*. 2014).

We performed hierarchical clustering among transcripts and samples using the limma-voom transformed log_2_(CPM) values through the ‘heatmap.2’ function in the ‘gplots’ package v.2.17.0 (Warnes *et al*. 2016). For each life-stage, samples were clustered based on Euclidean distance among transcript abundances and plotted on a heatmap by re-ordering the values by transcripts (rows) and by samples (columns). We evaluated the robustness of each cluster of transcripts identified through this method using the R package ‘fpc’ v.2.1.10 (Flexible Procedures for Clustering, Hennig 2015), which implements a bootstrapping algorithm on values of the Jaccard index to return a cluster stability index (Hébert 2016a). We only considered clusters with a stability index greater than 0.50 with 1,000 bootstraps for downstream Gene Ontology (GO) enrichment analyses. In total, 12 out of 16 clusters distributed across the two heatmaps were kept for GO enrichment analysis. We considered each cluster satisfying the stability index threshold as a module of co-expressed genes potentially bearing a broad functional status in accordance with the results from the GO enrichment analysis.

We identified functional categories over-represented in each co-expression module to characterize the biological functions associated with each life-stage. We used the Python package ‘goatools’ (Klopfenstein *et al*. 2015) to perform Fisher’s exact tests on GO annotation terms found in clusters of significantly differentially expressed gene. Annotation of GO terms for each gene was based on the published transcriptome of *Schistocephalus solidus* (Hébert *et al*. 2016b). GO terms over-represented in a given module, as compared to the reference transcriptome (FDR ≤ 0.05), were labelled as putative ‘transition-specific’ biological functions.

### Ecological annotation

We assigned an ecological annotation to transcripts exhibiting significant abundance changes between life stages or showing stage-specific expression patterns (Pavey *et al*. 2012). Two different types of ecological annotation were added to the dataset. First, we labelled un-annotated transcripts according to their significant variation in abundance across life stages. Information on GO terms over-represented in the cluster in which these transcripts could be found was also added. Second, we labelled transcripts showing “on-off patterns of expression” among stages and hosts as “stage-specific” or “host-specific”. A pre-defined specificity threshold was chosen as the log_2_(CPM) value representing the 5^th^ percentile of the distribution of the log_2_(CPM) across all transcripts. We identified stage-specific transcripts based on an average log_2_(CPM) above our pre-defined specificity threshold (‘ON’) across at least two-thirds of the worms in only one of the three life-stages. Similarly, we considered transcripts as host-specific if they showed an average log_2_(CPM) above the specificity threshold across two-thirds of the worms in any of the two hosts.

## RESULTS AND DISCUSSION

### Host transition as the main driver of genome reprogramming

A total of 2894 genes (28% of transcriptome) are significantly differentially regulated (FDR < 0.001) over the course of the infection of the fish and bird hosts (Tables S1). A multidimensional scaling analysis (MDS) performed on the top 1000 most differentiated genes in the dataset further suggests that the main factor that drives the divergence among individual worms is host type (fish vs. bird-gut model; Figure 1b). The first dimension of the MDS plot shows two distinct clusters: one with adult worms and another regrouping pre-infective and infective worms. This analysis also shows the grouping of pre-infective and infective worms into two different clusters on the second dimension. The distance on the first dimension between host types is at least twice as large as the distance separating pre-infective and infective worms, suggesting that host type is the main driving factor. This may largely be explained by physiological acclimatisation of the parasite to highly divergent thermal environments offered by the two hosts, or to other differences such as oxygen tension, pH or osmotic pressure (Smyth 1950; Aly *et al*. 2009; Oshima *et al*. 2011). The switch between these two hosts also correlates with rapid sexual maturation, reproduction and changes in energy metabolism (Clarke 1954). Altogether, these factors contribute to a major reprogramming of the worm transcription profile between hosts.

#### Biological activities focused towards reproductive functions

The development of the adult stage in the avian host requires the parasite to shift most of its biological activities from growth and immune evasion (Hopkins & Smyth 1951; Hammerschmidt & Kurtz 2005) to reproduction and possibly starvation (Hopkins 1950; Smyth 1954). In accordance with these life-history changes, sexual maturation pathways and reproductive behaviours were dominant functions in the transcriptional signature of the final host-switch, as supported by GO terms significantly enriched in co-expression modules (Figure 2a, Table S1). The largest co-expression module identified in the transition from infective plerocercoid to adult contains a total of 769 genes significantly over-expressed in adult worms (Figure 2a, cluster 4). This module is enriched (FDR < 0.05) in biological processes related to reproductive functions such as spermatid nucleus differentiation (GO:0007289), sperm motility (GO:0030317), luteinizing hormone secretion (GO:0032275) and positive regulation of testosterone secretion (GO:2000845) (Table S1). Early studies on the life cycle of *S. solidus* suggested that once the worm reaches the final bird host, its energy is canalized into maturation and reproduction, including egg-laying (Hopkins & Smyth 1951; Clarke 1954). Adult worms sampled in this study were collected after five days of *in vitro* culture in a bird-gut model at 40°C, 3-4 days after the onset of gamete production (Smyth 1946; Smyth 1954). The transcriptional signature confirms this at the molecular level, as we have detected the induction of many genes involved in sperm motility and cilium movement (Table S1).

**Figure 2.**
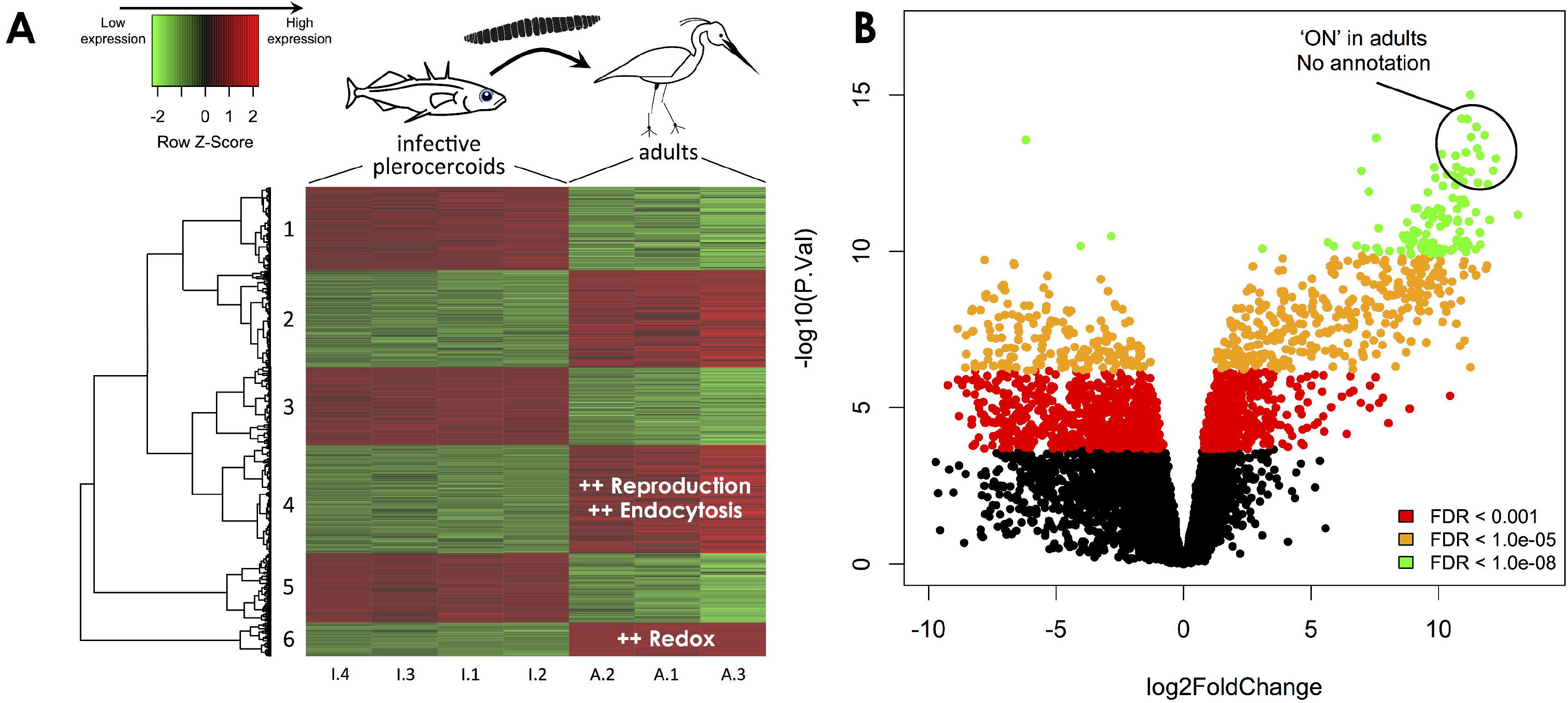
Differential patterns of gene expression reveal a strong stage-specific functional signature dominated by reproduction-associated activities in adult worms. A) Hierarchical cluster analysis showing co-expression relationships between genes significantly differentially expressed between infective plerocercoids and adult worms. Biological processes significantly enriched in each module appear in white on the heatmap. B) Volcano plot showing genes differentially expressed at three levels of FDR significance. Positive values of log2FoldChange correspond to up regulated genes in adult worms. Data points circled on the graph represent 18 of the top 30 most differentiated genes to which no annotation could be assigned. The un-annotated genes are turned ‘ON’ in adult worms and are part of the redox homeostasis functional module (heatmap cluster 6).

#### Re-organisation of the energy budget

The transition from infective plerocercoid to adult is characterized by a significant shift in energy metabolism (Barrett 1977). Empirical data suggests that during the first hours of maturation and reproduction, worms utilize glycogen reserves accumulated in the fish host (Hopkins 1950). Adult worms cultured *in vitro* are also capable of absorbing glucose after more than 40 hours at 40°C (Hopkins 1952), suggesting they can stop using their glycogen reserves and instead use host-derived nutrients. Our results suggest a complex and subtle pattern of regulation in terms of carbohydrate metabolism. In total, eleven steps of the glycolysis pathway were differentially regulated between the infective plerocercoid and adult stages (Figure 3). The first step in glycogen breakdown consists in converting glycogen to glucose-1-phosphate, a reaction catalysed by the enzyme glycogen phosphorylase (Smyth & McManus 2007). This enzyme is strongly up-regulated in adult worms (logFC = 5.9, FDR < 0.0001), suggesting an active use of glycogen reserves at this stage. The first three major biochemical transformations leading to glucose breakdown into more simple sugars are strongly down-regulated (Figure 3). Intriguingly, genes coding for the enzyme that produce glyceraldehyde-3-phosphate (GADP) are consistently up-regulated in adult worms. All three homologous genes identified as fructose-bisphosphate aldolase in our dataset, the enzyme responsible for the production of GADP, were labeled as being switched ‘ON’ in adults (see Materials and Methods for details). Most of the downstream genes leading to the production of pyruvate are down-regulated, with the exception of enolase, the enzyme responsible for the penultimate step of glycolysis, i.e. the conversion of glycerate-2-phosphate into phosphoenol-pyruvate (Figure 3, Table S1). Consistent with the semi-anaerobic conditions experienced by adult worms, we found a significant up-regulation of the gene coding for L-lactate dehydrogenase (logFC = 5.8, FDR < 0.001), the enzyme responsible for the conversion of pyruvate to lactate when oxygen supplies are low (Smyth & McManus 2007).

**Figure 3.**
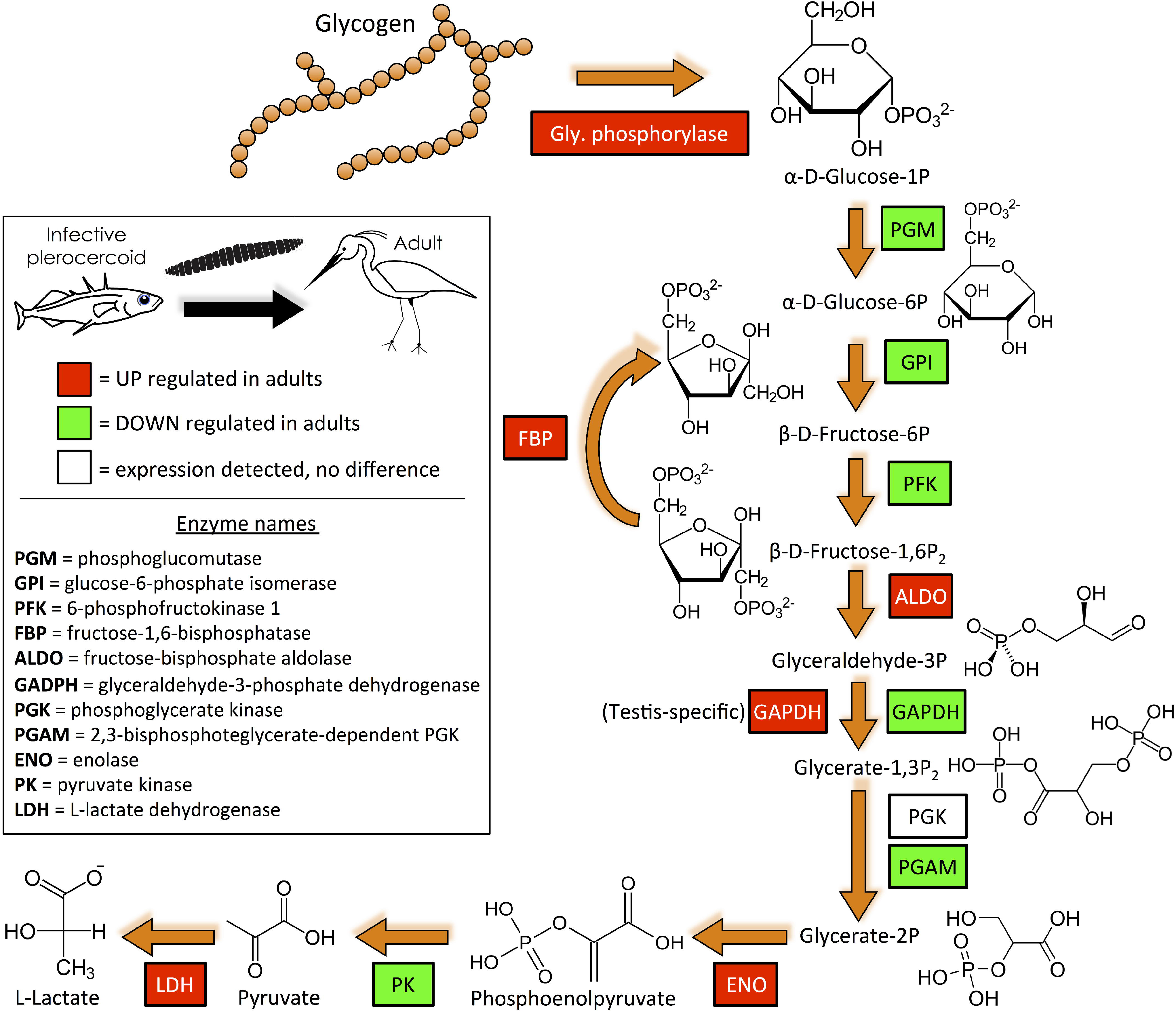
Partial glycolysis KEGG pathway highlighting the biochemical steps for which differential expression was detected between infective plerocercoids and adult worms. Boxes with a solid black line and white filling represent genes for which expression was detected with no significant difference between developmental stages. Red boxes represent up regulated genes in adult worms. Figure based on the complete KEGG pathway for glycolysis/gluconeogenesis (http://kegg.jp).

Interestingly, we found a testis-specific gene, coding for glyceraldehyde-3-phosphate dehydrogenase, among the few up-regulated genes of the glycolysis pathway. The gene was significantly over-expressed in adult worms, with a fold-change of 97 (logFC = 6.6, FDR < 0.0001). A homologous gene – with the same annotation, but not testis-specific – is conversely down-regulated in adults (logFC = -1.6, FDR = 0.0003). These results suggest that late stages of adult *S. solidus* may still be very active in terms of sperm production, even after several days in the avian host. At this late stage, the adult parasite may direct all of its energetic activities towards sperm production, in order to maximize rates of egg fertilization.

#### Potential role for endocytosis in balancing energetic reserves

How glucose is produced or acquired by adult *S. solidus* is unclear, but this activity could be performed by molecular mechanisms such as endocytosis or pinocytosis (Hopkins *et al*. 1978). This hypothesis led to the prediction that expression of genes specific to this pathway should be induced. Results from the GO enrichment analysis show a significant over-representation of biological processes related to endocytosis in adult worms. In total, 21 genes annotated with functional terms such as clathrin coat assembly (GO:0048268), clathrin-mediated endocytosis (GO:0072583), and regulation of endocytosis (GO:0006898) are co-regulated within the same cluster as reproduction-specific genes (Figure 2a, cluster 4). All 21 genes are significantly over-expressed in adult worms, as compared to the previous infective stage (Table S1). Even though we detect an over-expression of certain genes in adult worms that are involved in general mechanisms of endocytosis, we cannot determine where exactly these genes are expressed in the worm, since our experiment was performed on whole worms. They could be over-expressed in cells from the integumentary system, but also in other organs that are not involved in interaction with the external environment of the worm.

#### Regulation of redox pathways through novel species-specific genes

Adult stages of cestodes like *S. solidus* are exposed not only to the reactive oxygen species (ROS) produced by their own metabolism, but also to the ones generated by their host (Williams *et al*. 2013). Considering the extensive muscular activity required during reproductive behaviours (Smyth 1952; Clarke 1954) and the potential internalisation of host molecules by adult worms – which could include ROS produced by the host – maintenance of redox homeostasis should be a central activity performed at this stage. This scenario is reflected in the smallest co-expression module characterising the passage to the simulated avian host (Figure 2a, cluster 6), which harbours genes predominantly up-regulated in adults. This module does not exhibit significant enrichment for a particular biological activity, but it is nonetheless associated with oxidative stress and antioxidant metabolism, such as glutathione metabolic process (GO:0006749), glutathione biosynthetic process (GO:0006750) and glutathione dehydrogenase (ascorbate) activity (GO:0045174). Interestingly, of the 242 genes contained in this module, 174 (72%) are turned ‘ON’ in adults and ‘OFF’ in pre-infective and infective plerocercoids. All of the genes turned ‘ON’ in adult worms are found exclusively in this cluster (Table S1). Furthermore, 108 (62%) of these 174 ‘ON’ genes find no homology to any known sequence database, nucleotides or amino acids, while they are among the top differentiated genes in the final developmental transition (Figure 2b). This module is thus mainly composed of unknown genes that are co-expressed with oxidative stress genes being specifically up-regulated at the adult stage. Experimental evidence on the metabolism of adult worms shows a significant increase in lactate concentration at this stage (Beis & Barrett 1979), which is confirmed in our data by the increased expression of lactate dehydrogenase (Figure 3). Higher intracellular lactate content is considered as evidence for a more oxidised cytoplasm in mature worms (Beis & Barrett 1979). The redox module identified in our data supports this hypothesis of increased oxidative stress in adult worms, suggesting the importance of preventing the damage caused by ROS in late stages of infection.

### Detecting distinct developmental stages within the same host

#### Early plerocercoids associated with growth and regulatory programs

Evidence from physiological and morphological studies suggests that growth and organ development are the major biological programs that differentiate pre-infective from infective plerocercoids (Clarke 1954). *In vitro* experiments showed that in the first 48 hours following infection, the number of proglottids – i.e. body segments – is definitive. Unlike most of the cyclophyllidean tapeworms, *S. solidus* plerocercoids increase their bulk several hundredfold by adding layers of muscle tissue rather than adding proglottids (Hopkins & Smyth 1951; Clarke 1954). This suggests that organ development and tissue differentiation are switched off at this point, while muscle synthesis and growth are switched on. We examined if this developmental turning point is detectable in regulatory patterns of gene expression when comparing pre-infective versus infective plerocercoids.

Overall, three out of the four co-expression modules up-regulated in pre-infective plerocercoids were strongly associated with growth, cell division and regulatory functions. The first module contained 478 genes predominantly up-regulated in pre-infective plerocercoids (Figure 4a, cluster 6) and significantly enriched in GO terms related to DNA/RNA metabolism – e.g. replication, transcription and translation (Table S1). The second cluster contained 388 genes, also up-regulated in pre-infective plerocercoids (Figure 4a, cluster 4), and significantly enriched in biological activities involved in the regulation of cell cycle and cell division (Table S1). Among genes annotated with these GO terms, those exhibiting the largest expression difference between pre-infective and infective plerocercoids (i.e. logFC > 1.5, FDR < 0.001) code for mRNA splicing factors, DNA polymerase and key proteins involved in the regulation of mitosis (Table S1).

**Figure 4.**
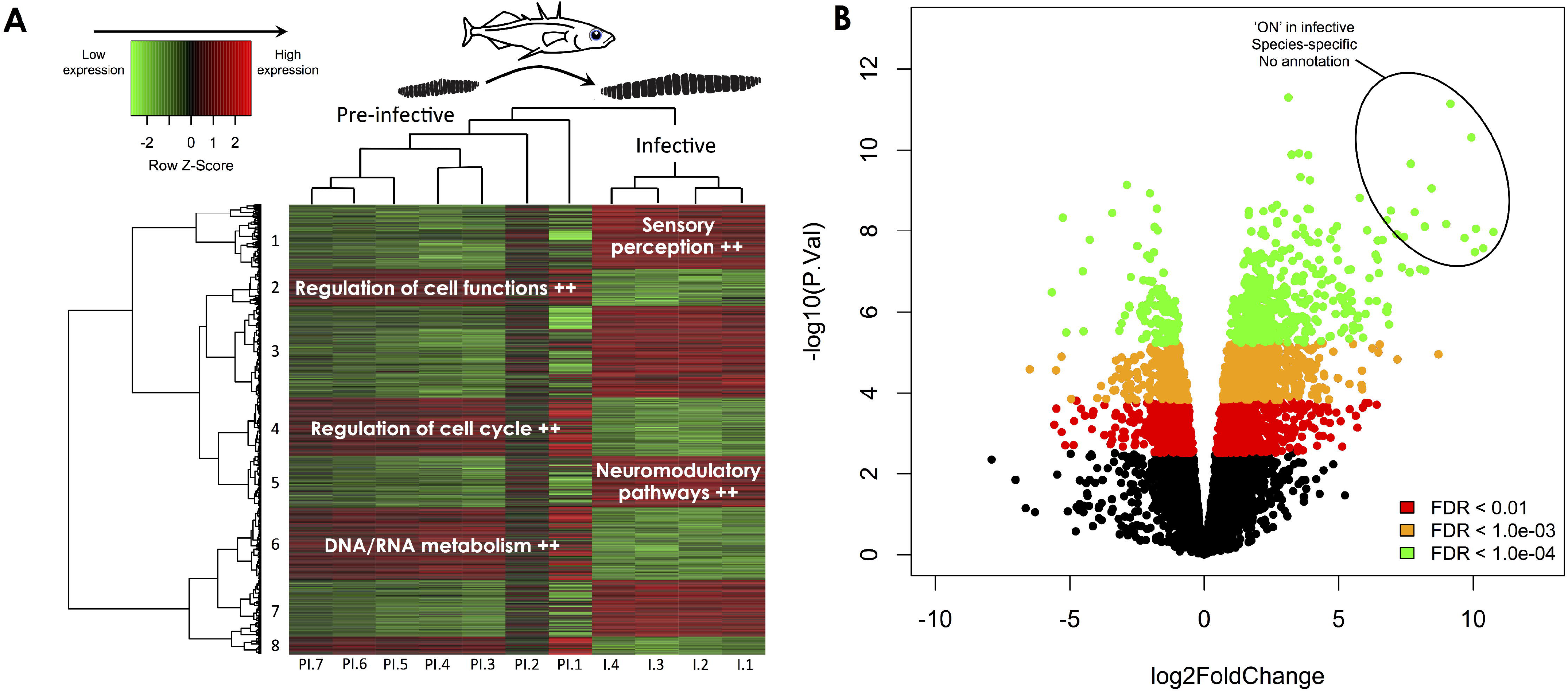
Differential patterns of gene expression suggest a significant role for neuromodulatory pathways in the development of infectivity towards the final host. A) Hierarchical cluster analysis showing co-expression relationships between genes significantly differentially expressed between pre-infective and infective plerocercoids. Biological processes significantly enriched in each module appear in white on the heatmap. B) Volcano plot showing genes differentially expressed at three levels of FDR significance. Positive values of log2FoldChange correspond to up regulated genes in infective plerocercoids. Data points circled on the graph represent the top 15 most differentiated genes, of which 100% are un-annotated. These uncharacterised sequences are all species-specific and systematically turned ‘ON’ in infective worms.

Developmental trajectories involving mitotic replications, cellular growth and tissue differentiation are often associated with specific regulatory processes that coordinate the timing of these events. Our results show that a third co-expression module significantly enriched in regulatory activities may perform this task. The module of 240 genes significantly up-regulated in pre-infective plerocercoids (Figure 4a, cluster 2) contains 144 (60%) genes annotated with GO terms. Among these, 17 genes (12% of annotated genes) have enriched GO terms related to regulatory processes involved in cellular functions such as apoptosis, mitosis, phosphorylation and cell division (Table S1). The genes with the largest difference in expression level are the transcription factors Sox-19b and GATA-3, with logFC values of 3.0 and 2.6 respectively (FDR < 0.001). Other regulatory genes include WNT4 and WNTG, with logFC values of 1.7 and 1.6 respectively (FDR < 0.001). Interestingly, these 17 regulatory genes are co-regulated, within this module, with other genes associated to cell cycle and DNA/RNA metabolism. In total, 41 genes (28% of annotated genes in the cluster) have enriched GO terms related to biological activities such as DNA replication, cell cycle, and DNA biosynthetic processes. These results suggest that small pre-infective plerocercoids activate a series of regulatory pathways in their intermediate fish host. We propose that these regulatory changes could result in the rapid increase in overall body mass through tissue differentiation, muscle fibre synthesis, organ formation and increased organ size (Benesh *et al*. 2013).

#### Specific transcriptional signature of infectivity dominated by environment sensing and un-annotated genes

One of the proxies used to infer infectivity in *S. solidus* plerocercoids is its significant influence on the immune system and behaviour of its fish host (reviewed in Hammerschmidt & Kurtz 2009), which implies communication between the two species (Adamo 2013). Consistent with this, we find that environmental sensing is the dominant function represented in the transcriptional signature of infectivity. The most compelling evidence comes from a large module of co-expressed genes significantly up-regulated in infective plerocercoids (Figure 4a, cluster 1). This module contains 407 genes significantly enriched in biological activities related to the cellular response of the organism to various molecules from the external and internal environment. More specifically, 84% of the 70 enriched GO terms in the module are involved in cellular responses to drugs and neuromodulators, and secretion and transport of various molecules through the cell membrane (Table S1). Of the 157 genes with a GO annotation in this module, 28 (18%) have GO terms involved in environmental sensing and interactions. Among these, those that exhibit the largest expression differences between pre-infective and infective plerocercoids code for proteins including monocarboxylate transporter 7 (logFC = 4.9, FDR = 2.2e-06), solute carrier family 22 member 21 (logFC = 4.6, FDR = 1.2e-05), multidrug resistance protein 1A (logFC = 4.4, FDR < 0.001), multidrug and toxin intrusion protein 1 (logFC = 4.4, FDR = 0.0013) and neuropeptide FF receptor 2 (logFC = 2.4, FDR = 3.6e-05).

The cluster described above is particularly interesting because of the high proportion of genes coding for unknown proteins differentially regulated between pre-infective and infective plerocercoids. One of the key features of this module is that GO annotations could be assigned to only 39% of the genes; hence it is the least annotated of all the modules characterizing the transition of plerocercoids from pre-infective to infective. The top 15 most differentiated genes between pre-infective and infective plerocercoids – with logFCs of 8-11, and FDRs < 0.00001 – are all completely unknown (Figure 4b), and are all *S*. solidus-specific sequences, i.e. we find no homology match to any known database except the *S. solidus* genome. These sequences were previously identified as species-specific transcripts based on a sequence homology analysis of the reference transcriptome (Hebert et al. 2016b). The transcriptome of *S. solidus* was compared to seven other parasitic transcriptomes (*B. malayi, C. sinensis, H. microstoma, T. solium, E. granulosus, E. multilocularis, S. erinaceieuropaei*) in a phylogenetic analysis. Results showed that 1 637 genes, which account for 19% of all protein coding genes, were *S*. solidus-specific (Hébert et al. 2016b). These sequences also exhibit valid open reading frames and are all highly expressed only in infective worms – i.e. they are turned ‘OFF’ in pre-infective plerocercoids and adult worms. The only information that can be used to assign a preliminary function to these genes is the ecological annotation that stems from our transcriptomic analysis (see Materials and Methods for details). These sequences were thus labelled as infective-specific and co-expressed with genes involved in environmental sensing and interaction (table S1). They might hold important, yet hidden, functional aspects that would allow a complete understanding of the interaction between infective plerocercoids and their fish host (Koziol *et al*. 2016).

#### Regulation of neural pathways could be essential for successful transmission

Our findings regarding the strong signal detected for neural pathways, such as environmental sensing, are further supported by another co-expression module. This module contains a total of 335 genes significantly up-regulated in infective worms (Figure 4a, cluster 5), among which 70% have GO annotations. Our results indicate that 41 (17%) of annotated genes in the module have GO annotations enriched in activities performed by the nervous system, while 124 (53%) of them have GO annotations related to transmembrane structure and activity. Biological processes associated with these genes include signal transduction, synaptic transmission, sensory receptor activities and synaptic exocytosis (Table S1). An interesting candidate emerges as one of the top differentiated genes in the module, with an expression fold change of 3.3 (FDR = 0.0003) between pre-infective and infective plerocercoids. This candidate is 5-hydroxytryptamine A1-alpha receptor, a serotonin receptor. Serotonin is an important regulator of carbohydrate metabolism, host-parasite communication and rhythmical movements – in conjunction with other related bioamines – in several cestodes (Marr & Muller 1995). Our results show a systematic up-regulation of serotonin receptors, adenylate cyclase and sodium-dependent serotonin transporters (SC6A4) specifically in infective worms (Figure 5). Functional studies showed that the signalling cascade of serotonin stimulates muscle contraction and glycogen breakdown in *Fasciota hepatica* and *Schistosoma mansoni* (Marr & Muller 1995). The ultimate downstream effect of serotonin signalling would be a cellular response to catabolize glycogen, the main source of energy in cestodes. In *S. mansoni*, it has been suggested that the main source of 5-HT is the host, even though some of the enzymes involved in the process and recycling of 5-HT have been detected in this species (Marr & Muller 1995). This is also the case with our dataset, in which we find at least one enzyme that is capable of breaking down one of the metabolites required for serotonin biosynthesis, i.e. tryptophan – indoleamine 2,3-dioxygenase 2, up regulated in infective worms with logFC = -3.1 and FDR = 0.01. According to the current transcriptome annotation (Hébert *et al*. 2016b), there is no sequence in the transcriptome of the pre-infective, infective and adult stages that is annotated as part of the biosynthetic process of serotonin. If serotonin metabolism plays such a central role in the success of *S. solidus* in its fish host without being synthesized by the worm itself, we could consider the possibility that it progressively uses the host’s supplies as it grows.

**Figure 5.**
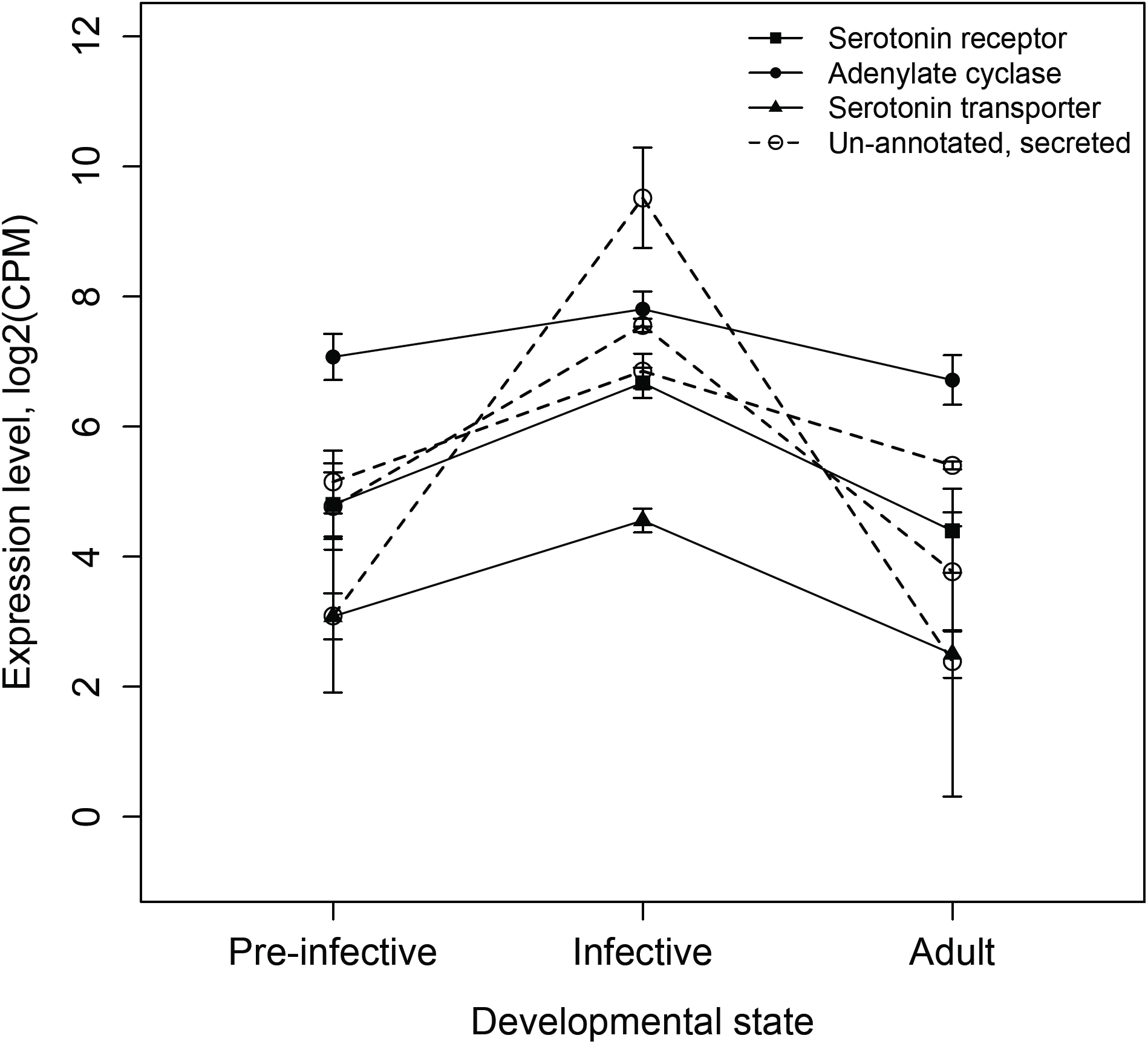
Activation of serotonin-related genes in the transcriptional signature of infectivity. Each data point on the graph corresponds to the average gene expression level (log2CPM) at a given developmental stage. Vertical bars represent the 95% confidence interval of the geometric mean. Genes coding for serotonin receptors (5-HT1A), adenylate cyclase (AC) and sodium-dependent serotonin transporters (SC6A4) are up-regulated specifically in infective plerocercoids. Un-annotated genes co-expressed in the same modules as serotonin-related genes and enriched in biological processes related to synaptic transmission and neural pathways show very similar patterns of expression (open circles with dashed lines). These un-annotated genes were labeled as ‘secreted’ based on the presence of a signal peptide in their sequence.

Successful completion of a complex life cycle involves intricate interactions between the parasite’s developmental program and physiological parameters experienced in each host. Investigating the transcriptomic signature of each developmental stage has led to the discovery of multiple novel yet un-annotated transcripts. These transcripts hold significant co-regulatory relationships with environmental interaction genes. Future functional characterization of these parasite-specific sequences promise to reveal crucial insights on how developmental and infection mechanisms evolved in different parasitic taxa.

## ACKNOWLEDGEMENTS

The authors would like to thank BJ Sutherland for comments and discussion, and M. Caouette and C. Tiley for precious technical work. This work was funded by a FRQ-NT grant to NAH and CRL, a Natural Science and Engineering Research Council of Canada (NSERC) Discovery grant to NAH, a NSERC Vanier Canada Graduate Scholarship and a Ressources Aquatiques Québec (RAQ) International internship fellowship to FOH, and a UK BBSRC MITBP fellowship to SG. CRL holds the Canada Research Chair in Evolutionary Cell and Systems Biology.

## ETHICS

Fish were captured under UK Environment Agency permit and with the permission of the landowner. All experiments were undertaken under a UK Home Office license (PPL80/2327) held by IB, in accordance with local and national regulations and with ABS/ASAB guidelines for the ethical treatment of animals in behavioral research (available online at http://asab.nottingham.ac.uk/ethics/guidelines.php).

## DATA ACCESSIBILITY

- Raw sequencing data: Sequence Read Archive (SRA) accession number SAMN04296611
- Final transcriptome sequences: NCBI BioProject PRJNA304161, uploaded with annotation.
- All data associated with the transcriptome: http://gigadb.org/dataset/100197
- Protocols: *In vitro/in vivo* culturing techniques and protocols available via protocols.io (doi: 10.17504/protocols.io.ew9bfh6).
- Python and R codes used for data analysis are available through github (Hébert 2016a,b).

## AUTHORS’ CONTRIBUTIONS

FOH, IB, CRL and NAH conceived the study. FOH and SG did the laboratory infections. FOH, SG and IB undertook fish dissections and parasite culture. FOH extracted RNA, prepared the sequencing libraries, performed bioinformatic analyses with supervision from NAH and CRL. FOH, CRL and NAH drafted the manuscript with input from IB. All authors read and approved the final manuscript.

## SUPPORTING INFORMATION

Table S1 Integrated results from the differential gene expression analysis for the complete transcriptome of *Schistocephalus solidus*. Summarises the information pertaining to the results form all analyses for each transcript included in the transcriptome, including unigene ID, gene product, GO annotation, association with co-expression modules, ON/OFF status, and differences in expression levels for each life stage transition (logFC with corresponding FDR value).

## COMPETING INTERESTS

The authors declare that they have no competing interests.

## REFERENCES

Adamo SA (2013) Parasites: evolution’s neurobiologists. The Journal of experimental biology, 216, 3–10.

Aly ASI, Vaughan AM, Kappe SHI (2009) Malaria parasite development in the mosquito and infection of the mammalian host. Annual Review of Microbiology, 63, 195–221.

Auld SK, Tinsley MC (2014) The evolutionary ecology of complex lifecycle parasites: linking phenomena with mechanisms. Heredity, 114, 125–132.

Barber I (2013) Sticklebacks as model hosts in ecological and evolutionary parasitology. Trends in Parasitology, 29, 556–566.

Barber I, Scharsack JP (2010) The three-spined *stickleback-Schistocephalus solidus* system: an experimental model for investigating host-parasite interactions in fish. Parasitology, 137, 411.

Barber I, Svensson PA (2003) Effects of experimental *Schistocephalus solidus* infections on growth, morphology and sexual development of female three-spined sticklebacks, *Gasterosteus aculeatus*. Parasitology, 126, 359–367.

Barber I, Walker P, Svensson PA (2004) Behavioural Responses to Simulated Avian Predation in Female Three Spined Sticklebacks: The Effect of Experimental *Schistocephalus solidus* Infections. Behaviour, 141, 1425–1440.

Barrett J (1977) Energy metabolism and infection in helminths. Symposia of the British Society for Parasitology, 15, 121–144.

Beis I, Barrett J (1979) The contents of adenine nucleotides and glycolytic and tricarboxylic acid cycle intermediates in activated and non-activated plerocercoids of *Schistocephalus solidus* (Cestoda: Pseudophyllidea) International journal for parasitology, 9, 465–468.

Benesh DP, Chubb JC, Parker GA (2013) Complex life cycles: why refrain from growth before reproduction in the adult niche? The American Naturalist, 181, 39–51.

Charles GH, Orr TS (1968) Comparative fine structure of outer tegument of *Ligula intestinalis* and *Schistocephalus solidus*. Experimental Parasitology, 22, 137–149.

Clarke AS (1954) Studies on the life cycle of the pseudophyllidean cestode *Schistocephalus solidus*. Proceedings of the Zoological Society of London, 124, 257–302.

Conradt U, Peters W (1989) Investigations on the occurrence of pinocytosis in the tegument of *Schistocephalus solidus*. Parasitology Research, 75, 630–635.

Davidson NM, Oshlack A (2014) Corset: enabling differential gene expression analysis for *de novo* assembled transcriptomes. Genome biology, 15, 1–14.

Hammerschmidt K, Kurtz J (2005) Surface carbohydrate composition of a tapeworm in its consecutive intermediate hosts: individual variation and fitness consequences. International journal for parasitology, 35, 1499–1507.

Hammerschmidt K, Kurtz J (2007) *Schistocephalus solidus:* Establishment of tapeworms in sticklebacks – fast food or fast lane? Experimental Parasitology, 116, 142–149.

Hammerschmidt K, Kurtz J (2009) Ecological immunology of a tapeworms’ interaction with its two consecutive hosts. In: Advances in Parasitology Advances in Parasitology. pp. 111–137. Elsevier.

Hébert FO (2016a) Bulk codes for RNA-seq analysis. Zenodo.

Hébert FO (2016b) corset_pipeline: First complete release. Github.

Hébert FO, Grambauer S, Barber I, Landry CR (2016a) Protocols for “Transcriptome sequences spanning key developmental states as a resource for the study of the cestode *Schistocephalus solidus*, a threespine stickleback parasite.” Protocols.io. (doi:10.17504/protocols.io.ew9bfh6)

Hébert FO, Grambauer S, Barber I, Landry CR, Aubin-Horth N (2016b) Transcriptome sequences spanning key developmental states as a resource for the study of the cestode *Schistocephalus solidus*, a threespine stickleback parasite. GigaScience, 5, 24.

Hébert FO, Grambauer S, Barber I, Landry CR, Aubin-Horth N (2016c) Resource for “Transcriptome sequences spanning key developmental states as a resource for the study of the cestode *Schistocephalus solidus*, a threespine stickleback parasite.” GigaScience Database. (doi:10.5524/100197)

Hopkins CA (1950) Studies on cestode metabolism. I. Glycogen metabolism in *Schistocephalus solidus* in vivo. Journal of Parasitology, 36, 384–390.

Hopkins CA (1952) Studies on cestode metabolism. II. The utilization of glycogen by *Schistocephalus solidus* in vitro. Experimental Parasitology, 1, 196–213.

Hopkins CA, Smyth JD (1951) Notes on the morphology and life history of *Schistocephalus solidus* (Cestoda: Diphyllobothriidae. Parasitology, 41, 283–291.

Hopkins CA, Law LM, Threadgold LT (1978) *Schistocephalus solidus:* pinocytosis by the plerocercoid tegument. Experimental Parasitology, 44, 161–172.

Klopfenstein D, Pedersen B, Flick P et al. (2015) GOATOOLS: Tools for Gene Ontology. Zenodo. https://zenodo.org/record/31628 (doi:10.5281/zenodo.31628)

Koziol U, Koziol M, Preza M et al. (2016) *De novo* discovery of neuropeptides in the genomes of parasitic flatworms using a novel comparative approach. International journal for parasitology, 1–13.

Körting W, Barrett J (1977) Carbohydrate catabolism in the plerocercoids of *Schistocephalus solidus* (Cestoda: Pseudophyllidea). International journal for parasitology, 7, 411–417.

Langmead B, Salzberg SL (2012) Fast gapped-read alignment with Bowtie 2. Nature Methods, 9, 357–359.

Law CW, Chen Y, Shi W, Smyth GK (2014) voom: Precision weights unlock linear model analysis tools for RNA-seq read counts. Genome biology, 15, R29.

Lee DL (1967) The Structure and Composition of the Helminth Cuticle. In: Advances in Parasitology Volume 4 Advances in Parasitology. pp. 187–254. Elsevier.

Marr J, Muller M (1995) Biochemistry and Molecular Biology of Parasites. Academic Press, London.

Oshima K, Ishii Y, Kakizawa S et al. (2011) Dramatic transcriptional changes in an intracellular parasite enable host switching between plant and insect. Plos One, 6, e23242.

Pavey SA, Bernatchez L, Aubin-Horth N, Landry CR (2012) What is needed for next-generation ecological and evolutionary genomics? Trends in Ecology & Evolution, 27, 673–678.

Poulin R (2011) Evolutionary Ecology of Parasites. Princeton University Press, Princeton.

Robinson MD, McCarthy DJ, Smyth GK (2010) edgeR: a Bioconductor package for differential expression analysis of digital gene expression data. Bioinformatics, 26, 139–140.

Scharsack JP, Gossens A, Franke F, Kurtz J (2013) Excretory products of the cestode, *Schistocephalus solidus*, modulate *in vitro* responses of leukocytes from its specific host, the three-spined stickleback (*Gasterosteus aculeatus*). Fish & Shellfish Immunology, 35, 1779–1787.

Scharsack JP, Koch K, Hammerschmidt K (2007) Who is in control of the stickleback immune system: interactions between *Schistocephalus solidus* and its specific vertebrate host. Proceedings Of The Royal Society B-Biological Sciences, 274, 3151–3158.

Schjørring S (2003) Schistocephalus solidus: a molecular test of premature gamete exchange for fertilization in the intermediate host *Gasterosteus aculeatus*. Experimental Parasitology, 103, 174–176.

Smyth DJ (1946) Studies on tapeworm physiology, the cultivation of *Schistocephalus solidus* in vitro. The Journal of experimental biology, 23, 47–70.

Smyth JD (1950) Studies on tapeworm physiology. V. Further observations on the maturation of *Schistocephalus solidus* (Diphyllobothriidae) under sterile conditions in vitro. The Journal of parasitology, 36, 371.

Smyth JD (1952) Studies on tapeworm physiology. VI. Effect of temperature on the maturation *in vitro* of *Schistocephalus solidus*. The Journal of experimental biology, 29, 304–309.

Smyth JD (1954) Studies on tapeworm physiology. VII. Fertilization of *Schistocephalus solidus* in vitro. Experimental Parasitology, 3, 64–67.

Smyth JD, McManus DP (2007) The Physiology and Biochemistry of Cestodes. Cambridge University Press, Cambridge.

Threadgold LT, Hopkins CA (1981) *Schistocephalus solidus* and *Ligula intestinalis*: pinocytosis by the tegument. Experimental Parasitology, 51, 444–456.

Tierney JF, Crompton DW (1992) Infectivity of plerocercoids of *Schistocephalus solidus* (Cestoda: Ligulidae) and fecundity of the adults in an experimental definitive host, *Gallus gallus*. J Parasitol, 78, 1049–1054.

Warnes GR, Bolker B, Bonebakker L et al. (2016) gplots: Various R programming tools for plotting data. R package version 2.0.1.

Wilbur HM (1980) Complex life cycles. Annual review of Ecology and Systematics, 11, 67–93.

Williams DL, Bonilla M, Gladyshev VN, Salinas G (2013) Thioredoxin glutathione reductase-dependent redox networks in platyhelminth parasites. Antioxidants & redox signaling, 19, 735–745.

